# Rule Activation and Ventromedial Prefrontal Engagement Support Accurate Stopping in Self-Paced Learning

**DOI:** 10.1101/169110

**Authors:** Sean R. O’Bryan, Eric Walden, Michael J. Serra, Tyler Davis

## Abstract

When weighing evidence for a decision, individuals are continually faced with the choice of whether to gather more information or act on what has already been learned. The present experiment employed a self-paced category learning task and fMRI to examine the neural mechanisms underlying stopping of information search and how they contribute to choice accuracy. Participants learned to classify triads of face, object, and scene cues into one of two categories using a rule based on one of the stimulus dimensions. After each trial, participants were given the option to explicitly solve the rule or continue learning. Representational similarity analysis (RSA) was used to examine activation of rule-relevant information on trials leading up to a decision to solve the rule. We found that activation of rule-relevant information increased leading up to participants’ stopping decisions. Stopping was associated with widespread activation that included medial prefrontal cortex and visual association areas. Engagement of ventromedial prefrontal cortex (vmPFC) was associated with accurate stopping, and activation in this region was functionally coupled with signal in dorsolateral prefrontal cortex (dlPFC). Results suggest that activating rule information when deciding whether to stop an information search increases choice accuracy, and that the response profile of vmPFC during such decisions may provide an index of effective learning.

## 1. Introduction

Should I keep studying for my math test? Do I know enough about cars to pick out a good one? When gathering evidence for a decision, individuals are continually faced with the question: Have I learned enough yet? Learners must strike a compromise between collecting enough information to make accurate decisions while avoiding collecting redundant information and—consequently—wasting time and resources.

Currently, little is known about the neurobiological mechanisms that govern decisions about when to stop gathering new information. In the domain of value-based decision making, behavioral research has often focused on heuristics or stopping rules, such as take-the-best, that people employ when presented with cues of varying predictive value (Gigerenzer & Goldstein, 1996). The use of such strategies, however, can vary across participants, even in decision environments that encourage the use of a particular heuristic (Newell & Shanks, 2003; Newell et al., 2004). Thus recent research has begun to focus on participants’ use of confidence thresholds for determining when stopping is appropriate, as opposed to application of specific rules per se (Svenson, 1992; Karelaia, 2006; Hausmann & Läge, 2008). In the present study, we test the neural mechanisms that contribute to stopping decisions during learning, and how activation of information associated with a choice evolves leading up to when a decision threshold is reached.

Neurobiologically, in a recent study that required participants to take or decline sequentially-presented stock options, stopping of information search was found to engage anterior cingulate, insula, and ventral striatum (Costa & Averbeck, 2015). Additionally, accumulated value and reward associated with stopping decisions in sequential sampling paradigms have been associated with activation in lateral orbitofrontal cortex, vmPFC, and the basal ganglia (Gluth, Rieskamp, & Büchel, 2012; Costa & Averbeck, 2015). Although these results have shed light on the neural correlates of stopping in value-based choice, how they translate to stopping in learning contexts, such as when people make decisions about their mastery of new concepts, remains an open question.

Rule-based category learning provides an ideal context to examine the neural basis of stopping in learning because many real world concepts are associated with rules, and because the neural systems that support rule-based categorization are well-understood (for review, see Seger & Miller, 2010; Ashby & Maddox, 2011). Cognitively, rule-based category learning involves using hypothesis testing and selective attention to establish and focus on stimulus dimensions that are relevant for predicting category membership (Maddox & Ing, 2005; Zeithamova & Maddox, 2006). For example, when learning to distinguish between birds and mammals, people may learn to selectively attend to whether an organism has wings. Selective attention emerges over the course of learning to minimize prediction error, and has the effect of expanding the representation of dimensions that lead to successful categorization (Nosofsky, 1986; Kruschke, 1992; Folstein, Palmeri, & Gauthier, 2013). Neurobiologically, rule-based category learning is thought to depend on executive cortico-striatal loops connecting the prefrontal cortex with the head of the caudate (Seger & Miller, 2010), with the ventral striatum playing a particularly important role in both initial rule acquisition and reversal learning in the event of a rule switch (Seger & Cincotta, 2006; Liu et al., 2015). The medial temporal lobes are thought to be involved in the long-term maintenance and retrieval of these category rules (Poldrack et al., 2001; Davis, Love, & Preston, 2012).

Currently, how the neurobiological systems involved in rule-based category learning contribute to decisions to stop learning is not clear because the vast majority of neuroimaging studies have employed fixed numbers of trials and not given participants leeway in decisions about whether to continue learning. However, it is possible to infer what mechanisms may underlie decisions to stop learning by incorporating predictions from recent neuroimaging research on stopping decisions in value-based choice and research on confidence in choice behavior more generally. In value-based decision making, the vmPFC has been shown to track subjective value (Tom et al., 2007; Bartra et al., 2013), and is sensitive to cost-benefit discrepancies among response options (Basten et al., 2010; Lim, O’Doherty, & Rangel, 2011). These findings are consistent with results showing that the vmPFC tracks accumulated value in value-based stopping decisions. Recent findings suggest that vmPFC may also code general decision evidence or confidence associated with a choice, perhaps in parallel with value (Di Martino et al., 2013; Barron, Garvert, & Behrens, 2015; Lebreton et al., 2015). Indeed, a number of basic category learning tasks have found that the engagement of vmPFC is correlated with greater decision evidence for categorization choices (Grinbald et al., 2006; DeGutis & D’Esposito, 2007; Seger et al., 2015; Davis, Goldwater, & Giron, 2017). Thus we expect that decision evidence/confidence signals from the vmPFC may contribute to subjective thresholds participants use when deciding whether they have learned enough information.

In addition to the vmPFC, we also expect the dorsolateral PFC (dlPFC) to contribute to stopping decisions. Recent research has suggested that a region of the posterior dlPFC may be involved in comparing accumulated perceptual information to decision criteria when making perceptual decisions (Heekeren et al., 2004; White et al., 2012). In terms of decisions to stop gathering new information, the dlPFC may monitor information from a number of regions to determine when stopping criteria have been reached, including confidence signals from the vmPFC. Indeed, several studies have observed increased functional connectivity between the dlPFC and vmPFC when participants make decisions that require weighing the subjective values of different choice options (Baumgartner et al., 2011; Rudorf & Hare, 2014). A similar coupling between dlPFC and vmPFC may support computing decision criteria for stopping in category learning.

As a complement to univariate activation, which measures the extent to which brain regions are engaged leading up to decisions to stop, multi-voxel pattern analysis (MVPA) may provide an additional window into how participants are processing category information leading up to stopping decisions. Recent studies have found, as a result of learning, similarities between activation patterns elicited for members of a category come to reflect how participants attend to the stimulus dimensions, such that items sharing values along a rule dimension come to elicit more similar activation patterns (e.g., Mack, Preston, & Love, 2013; Mack, Love, & Preston, 2016). Such changes in neural similarity spaces may track participants’ decisions to stop learning. Specifically, we expect that as subjects selectively narrow their attention to a particular rule dimension leading up to a decision to stop gathering information, activation patterns associated with this dimension will become increasingly prominent in the underlying neural similarity space.

To test our predictions for engagement of the vmPFC/dlPFC and how rule-relevant information will be activated leading up to a stopping decision, we trained participants to categorize triads of visual stimuli using simple rules based on one of three binary stimulus dimensions (Figure 1). The stimulus dimensions consisted of three distinct visual categories: faces, objects and scenes. Participants learned, using trial and error, which stimulus dimension was the rule dimension and was predictive of category membership. Each full task trial was comprised of three to four fixation-separated subcomponents: a learning trial, which involved categorizing the visual stimulus; the presentation of correct/incorrect feedback; a decision trial where participants chose whether to solve the category rule or to continue learning; and finally, if a solve response was made, a trial for selecting between the three possible category rules. The use of a Type I category structure (Shepard, Hovland, & Jenkins, 1961) allowed participants to solve rules rapidly, allowing us to robustly measure how activation of rule-relevant information evolved over many different individual decisions to stop learning and solve the rule. To further maximize our ability to detect subtle changes in attentional weighting that result from learning, face, object, and scene images were used as stimulus dimensions. These real-world categories exhibit strong properties for representational decoding (Haxby et al., 2001; O’Toole et al., 2005), and were localized within each subject using independent scans to create ROIs that were unbiased to any potential learning effects (Davis et al., 2014). For the purposes of this study, the representational analysis was focused on the activation of patterns associated with the rule dimension that participants eventually chose as the solution during both the “learning” and “decision” portions of the trial (Figure 1).

**Figure 1.**
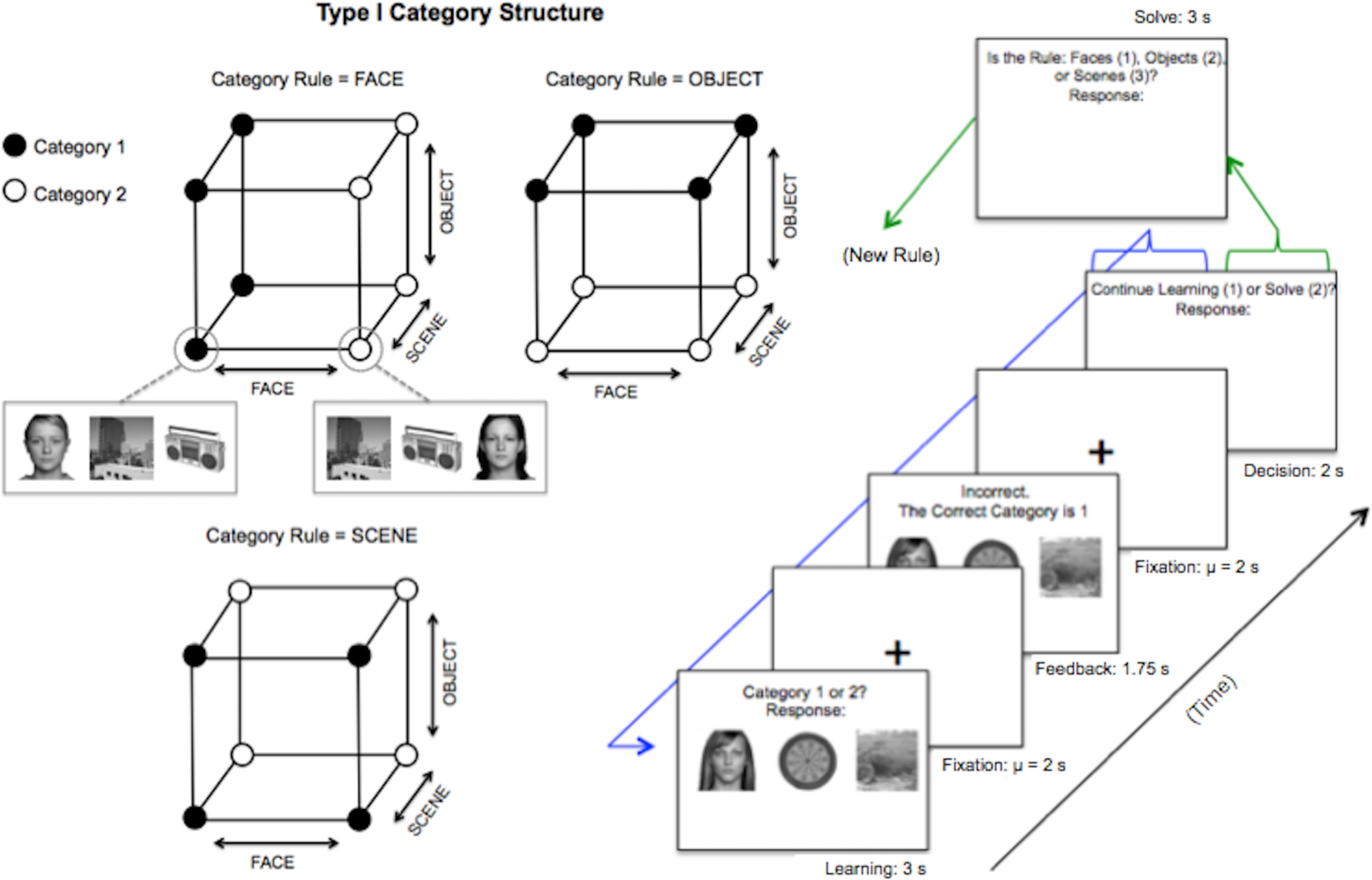
Abstract category structure and an example trial for the self-paced rule learning task. Any given rule follows a Type I category structure (Shepard, Hovland, & Jenkins, 1961), where only one stimulus dimension is the “rule dimension” and is predictive of the correct category during learning (e.g., if the category rule is Face: Face 1 = Category 1, Face 2 = Category 2). Each point on the cubes corresponds to one of eight possible exemplars (feature combinations) displayed on the screen. During the task, participants are prompted with the decision to solve the rule or continue learning after each response-feedback pairing. If ‘continue’ is chosen, a new feature combination is randomly selected from the same rule structure and the process repeats. If’solve’ is chosen, participants are taken to a screen where they are prompted to select the object class that constituted the rule dimension. Following a solve response, learning begins again with a random rule and new set of stimuli.

To successfully navigate our task, participants must first learn to selectively attend to the rule dimension to predict category membership. We hypothesized that the increasing selective attention to the rule dimension prior to stopping would result in multi-voxel patterns that become increasingly similar to the object class associated with the chosen rule dimension as participants’ neared a choice to stop. Once attention is allocated to the potential rule dimension, it is necessary for the participant to monitor the evidence consistent with this category rule. We predicted that vmPFC would be involved in representing the confidence or decision evidence for an attended rule dimension, with stopping marked by greater vmPFC activation than decisions to continue learning. Finally, participants must make a decision to stop learning once this evidence has reached a criterion. We hypothesized that dlPFC would track confidence/decision evidence signals from the vmPFC to determine whether a stopping threshold had been reached.

To preview our findings, the representational analysis revealed a dynamic neural accumulation process whereby activation of multi-voxel patterns associated with the object class eventually chosen as the rule dimension increased over the trials leading up to stopping decisions. Compared with the choice to continue learning, participants engaged a widespread network of brain regions including medial PFC when choosing to stop; within this contrast, activation of vmPFC and object-selective cortex were positively correlated with participants’ ability to solve rules accurately throughout the task. Moreover, we show that decisions to stop acquiring information and solve a rule are associated with enhanced functional connectivity between vmPFC and dlPFC.

## 2. Materials and Methods

### 2.1. Participants

Twenty-five healthy, right-handed volunteers (ages 21 – 57, mean ± SEM = 27.32 ± 1.67, 17 women) were recruited through online newsletters and flyers posted at Texas Tech University. All subjects were included in the final analysis. All subjects provided written informed consent prior to participation, and were compensated $35 for a 1.5 hour session. The study protocol was approved by the Texas Tech University Human Research Protection Program.

### 2.2. Experimental Paradigm

The experimental paradigm consisted of a self-paced category learning task completed over 5 consecutive fMRI scanning runs. Participants first learned to classify image triads as belonging to one of two categories based on feedback delivered following each learning trial. The three simultaneously presented features always consisted of one face, one object, and one scene. One object class was predictive of category membership and was the rule dimension. Subjects were instructed to learn the rule dimension that was predictive of category membership. Each dimension (object class) had two values, resulting in a category structure comprised of six total cues (Figure 1). The face, object, and scene images used were black-and-white squares presented on a white background with black text. The screen positions of each cue (left, right, or center) were randomized on each learning trial to avoid gaze effects. Within each scanning run, the stimuli used for each rule were drawn randomly from a set of 36 images for each respective class, and were removed from the set after they were used for a given rule. After each learning/feedback trial pairing, participants were given the option to explicitly solve the rule (indicating whether faces, objects, or scenes were predictive) or to continue learning. These decision trials were of primary interest to the present study, allowing us to isolate the neural and behavioral correlates of stopping.

Participants received instructions indicating that their goal was to correctly solve as many rules as possible over the course of the experiment. Beyond the overall instructions to focus on solving rules correctly, correct solves were not explicitly incentivized. Participants were provided with an on-screen summary of their performance (the number of rules solved correctly) at the end of each scanning run. Figure 1 displays a schematic of the self-paced category learning task. Procedurally, image triads were presented on the screen accompanied by the prompt “Category 1 or Category2?” and a response prompt, with a maximum response time of 3 s. Next, feedback was presented for 1.75 s, indicating whether the response was correct or incorrect accompanied by the correct response. Feedback included presentation of the associated stimulus triad to facilitate learning. Following feedback, participants were prompted to make a button press indicating whether they wished to continue learning or solve the rule. These decision trials had a 2 s response deadline, and did not present the associated image triad. If a continue response was given, the task proceeded to the next category learning trial for the current rule and stimulus set. If a solve response was given, participants were prompted to make a response as to whether the rule (predictive feature) was faces, objects, or scenes within 3 s. Regardless of whether a response was given in the allotted time, the rule and associated set of images were then randomly reset, and the task proceeded to a category learning trial for the new rule.

Variable fixation periods drawn from truncated exponentials (mean = 2 s) separated stimulus presentation from feedback, feedback from decision trials, decision from solve or the next category learning trial, and solve trials from the next category learning trial. The number of learning trials completed by each participant varied slightly due to the nature of the task and the necessity of a timed cutoff for each scanning run. The mean number of learning trials completed in the sample over 5 functional scans was 150.4 (Range = 140 – 159, *SD* = 4.28). To provide subjects with a performance index of their rule solving accuracy, at the conclusion of each scanning run a screen notified them of the number of rules they correctly solved during that scan. Participants received instructions and completed practice trials for the task outside of the scanner for approximately 10 minutes before engaging in the fMRI experiment.

Prior to beginning the rule learning task, participants completed two functional localizer scans. Each trial of the localizer task involved presenting a face, object, or scene individually on the screen, asking subjects to make a button press corresponding to the appropriate item category within 3 s. Variable fixation lengths drawn from a truncated exponential (mean = 2 s) separated each trial. Over the duration of the localizer phase, subjects categorized 38 examples of each object class. The black-and-white images used during the localizer runs were presented in a random order, and did not include any of the stimuli used for the experimental task.

### 2.3. Image Acquisition

Imaging data were acquired on a 3.0 T Siemens Skyra MRI scanner at the Texas Tech Neuroimaging Institute. MPRAGE anatomical scans provided high-resolution structural images of the whole brain in the sagittal plane for each participant (TR = 1.9 s; TE = 2.49 ms; *θ =* 9°; FoV = 240 x 240 mm; matrix = 256 x 256 mm; slice thickness = 0.9 mm, slices = 192). Functional images were acquired using a single-shot T2*-weighted gradient echo EPI sequence (TR = 2.09 s; TE = 25 ms; *θ =* 70°; FoV= 192 x 192 mm; matrix = 64 x 64; number of axial slices = 41, slice thickness = 2.5 mm; 0.5 mm gap), and slices were tilted to reduce orbitofrontal dropout (Deichmann et al., 2003).

### 2.4. Image Analysis and Preprocessing

Functional data were preprocessed and analyzed using FSL (www.fmrib.ox.ac.uk/fsl) and anatomical preprocessing was conducted with Freesurfer (autorecon1). Images were skull stripped, motion corrected, prewhitened, and high-pass filtered (cutoff: 60 s). For univariate analysis, data were spatially smoothed using a 6 mm FWHM Gaussian kernel. No spatial smoothing was used for representational similarity analysis. First-level statistical maps were registered to the Montreal Neurological Institute (MNI)-152 template using boundary-based registration (BBR) to align the functional image to the structural image, and 12 df to align the structural image to the MNI-152 template.

Three-level statistical analysis of the functional data was carried out using FEAT and FSL’s Randomise. At level-one, within-run associations between task regressors and functional time series were examined. Eight task regressors and their temporal derivatives were included in the level-one analysis, including the onsets of correct/incorrect learning trials, correct/incorrect feedback, decision trials receiving a continue response, decision trials receiving a solve response, and correct/incorrect rule-solving trials. Nuisance regressors included temporal derivatives for the task variables, trials in which subjects failed to make a response, realignment parameters from motion correction, their temporal derivatives, and volume-wise indicator variables for scrubbing volumes that exceeded a framewise displacement of 0.9 mm (Siegel et al., 2014). Task-based regressors were convolved with a double gamma hemodynamic response function, while motion parameters were left unconvolved. Prewhitening was performed at level-one to control for temporal autocorrelation in the hemodynamic response. At level-two, within-run parameter estimates for task variables were averaged across runs for each subject using a fixed effects model. At level-three, we averaged parameter estimates across subjects using a random effects model for population inference. *Z-*scored mean rule solving accuracy was included in the level-three model as a moderator to test the hypothesis of performance-based differences in neural engagement. Final statistical maps were corrected for multiple comparisons at p < .05 using a permutation-based cluster mass thresholding, implemented in FSL’s Randomise. This analysis included a primary (cluster-forming) threshold of *t* = 2.49 (critical value of *t* for df = 24 and alpha = 0.01), and 6 mm variance smoothing. Permutation-based tests are immune to recent concerns about potential inflation of family-wise error rates in parametric GRF-based cluster thresholding (Eklund, Nichols, & Knutsson, 2016).

### 2.5. Multivariate fMRI Analysis

To obtain trial-by-trial estimates of the hemodynamic response, we computed a β-map (Rissman, Gazzaley, & D’Esposito, 2004) for each stimulus onset using an LS-A procedure (Mumford et al., 2012), which involves modeling individual task trials as separate regressors in a single general linear model. The estimated neural activation patterns for each onset were then registered to standard space. The aim of the multi-voxel pattern analysis was to examine neural signatures indicating the processing of predictive versus non-predictive item classes over time leading up to a solve decision. To obtain featural selective attention predictions, we computed similarities between the patterns for each trial in the task and the patterns computed for face-, object-, and scene-only trials from the independent localizer. The β-series used to compute the multi-voxel patterns was spatially localized to a functional ROI spanning category-selective brain regions in ventral temporal/occipital cortex that included lateral occipital complex, fusiform gyrus, and parahippocampal cortex (Supplementary Figure A1). This mask was created by binarizing and combining the statistical maps for Object > Scene and Scene > Object from the independent localizer task. As opposed to creating three distinct ROIs for face-, object-, and scene-selective regions, the decision to use a larger cross-category ROI was motivated by the fact that measuring the relative neural similarity to each visual stimulus type at the trial level was crucial for the present analysis. In this scenario, using a combined mask allows pattern estimates for a given feature to vary according to neural activity in the regions specifically associated with that feature, while simultaneously accounting for pattern variability in the alternative regions. Given that attention to a certain object class should lead to high relative activation in the brain region that represents it and relatively low activation in regions that do not, the absence of activation in these alternative regions may add positive predictive value to similarity estimates. A Pearson correlation was used to compute correlation distances (1 – *r*) between trials in the category learning task with face, object, and scene patterns from the independent localizer task within each subject. Each subject-level correlation map was transformed using Fisher’s *Z* and aggregated over trial type for statistical comparison.

**Figure A1.**
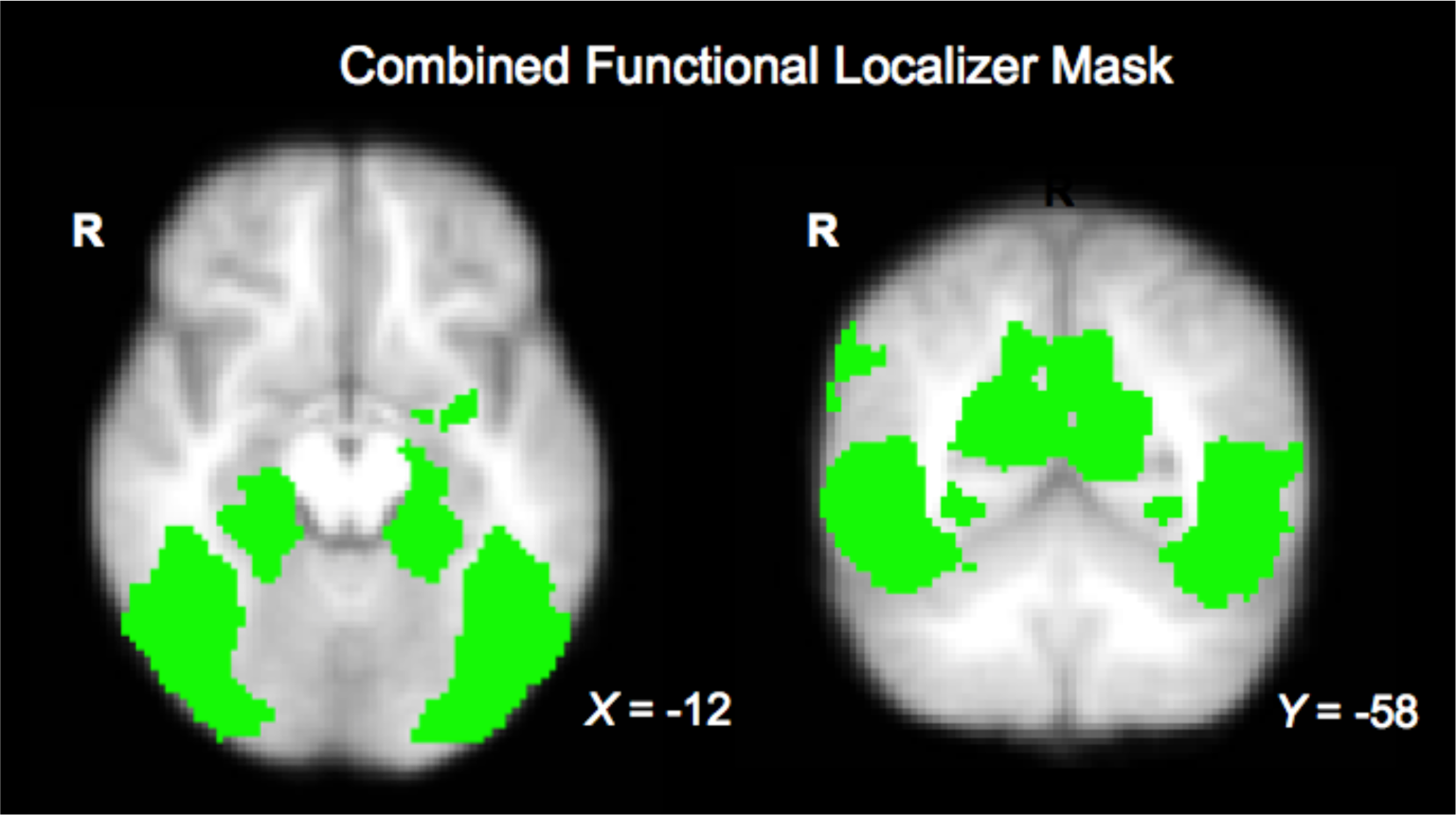
Functional localizer mask encompassing face-, object-, and scene-selective regions for the RSA learning analysis.

The primary goal of the multivariate analysis was to examine how temporal changes in the representation of rule information during learning support stopping decisions. Therefore, we first sorted learning trials according to the rules participants selected during the solve stage regardless of whether these solve responses were correct. We then counted back five trials from the solution trial to determine how the activation of information associated with the selected class changed as participants approached a solution. Five trials were analyzed in order to capture the average number of learning trials encountered by participants prior to solving (see Behavioral Results). For each learning trial, similarity estimates for the chosen object class were contrasted with the mean similarity to both of the unchosen object classes. To test whether there were significant time-based changes in neural similarity to the chosen item class during learning, we computed two multi-level models with similarity to chosen/unchosen as the outcome variables and 5-trial n-back to solve as a fixed predictor variable, allowing the intercept for each subject to vary randomly. Similarity output for the chosen/unchosen object classes over 3070 total observations served as the level-1 units for each model, nested within 25 participants. A random slope parameter for n-back was then added to each null model to examine whether the relationship between neural similarities and time differed between subjects. We determined whether to proceed with the inclusion of each random effect parameter (i.e., random time slopes and intercepts for each subject) by comparing AIC between model fits. To determine how the time-course of attention differed across instances of effective and ineffective learning, the same series of analyses were then conducted for similarity to the chosen object class for the learning trials preceding correct and incorrect solves, respectively.

### 2.6. vmPFC Signal Analysis

To assess how vmPFC signal may covary with the temporal distance from a solve decision, subject-level accuracy, and neural similarity to the chosen object class, we conducted two multi-level regression models with trial-by-trial vmPFC signal during the decision phase as the outcome variable. Both models included n-back and subject-level accuracy as fixed predictor variables, allowing subject intercepts to vary randomly. The two models differed only in that the first model included fixed predictors for similarity to the chosen and unchosen object classes during the immediately preceding learning trial, whereas the second model included predictors for similarity to the chosen/unchosen object classes from the decision trials. For these analyses, the signal in vmPFC was measured via mean activation estimates from the beta-series regression, masked anatomically within bilateral frontal medial cortex from the Harvard-Oxford Atlas.

### 2.7. PPI Analysis

A psychophysiological interaction analysis was conducted to examine whether any regions of the brain were functionally coupled with vmPFC during stopping decisions. The specific location of the vmPFC ROI (Figure 5A) was obtained by binarizing and anatomically masking the statistical map for Solve > Continue using bilateral frontal medial cortex from the Harvard-Oxford Atlas. FEAT 3-level analysis was then conducted, starting with the same eight level-1 explanatory variables used in the primary univariate model (see image analysis for further detail). Further, the model included *z-*scored time series for the vmPFC seed region, an interaction between Continue (centered) and the vmPFC time series (mean), and an interaction between Solve (centered) and the vmPFC time series (mean). The model included the same nuisance variables as listed above. The contrast of interest for the PPI was the difference between interactions: solve*vmPFC time series > continue*vmPFC time series. Final statistical maps were obtained using FSL’s Randomise and the same thresholding settings described in the image analysis section.

## 3. Results

### 3.1. Behavioral Results

The average number of learning trials encountered prior to a solve response and subsequent rule switch was 4.48, with mean learning trial accuracy at 57.2% (*SD* = 7.94%, range = 45.9% – 74.5%). Performance curves for category learning trials preceding both correct and incorrect rule solves are displayed in Figure 2. The mean number of solve responses (and thus, rules encountered) by subjects over the 5 task runs was highly variable (*M* = 33.6, *SD* = 10.5, range = 11 – 61), which was expected due to the free-responding nature of the task. Group performance for rule solving was well above chance (*M* = 66.1%, *SD* = 24.5%, range = 19.4% − 94.4%), although 7 participants failed to perform significantly above chance. Comparison of learning behavior across task performance revealed a significant negative correlation between the number of solve responses made throughout the task and mean rule solving accuracy (*r* = −.444, *t* (23) = − 2.37, *p* = .026), raising the possibility that some participants may have focused on completing a higher total volume of rules as opposed to focusing on solving fewer accurately. Similarly, the number of correct learning trials prior to solving the rule was positively associated with rule solving accuracy (*r* = .504, *t* (23) = 2.80, *p* = .010). In particular, subjects with below-chance rule solving accuracy chose to solve the rule after making fewer correct responses (*M* = 1.95) than those who solved rules with above-chance accuracy (*M* = 3.39).

**Figure 2.**
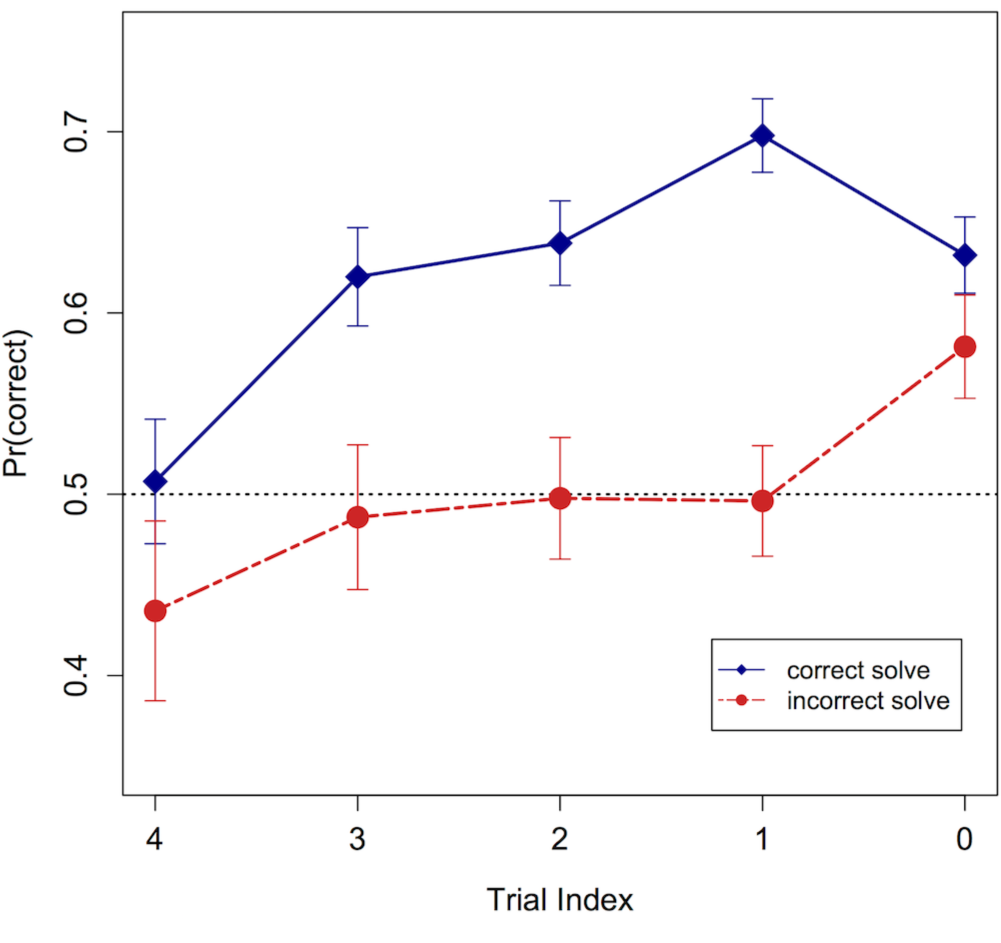
Learning performance curves preceding correct and incorrect solves. On the x-axis, 4:0 represent category learning trials preceding a solve decision, with zero being the final learning trial before participants opted to stop and solve the rule. Mean proportion correct is depicted on the y-axis. The horizontal line at 50% correct represents chance performance.

### 3.2. MVPA Results: Learning

To solve for a rule, participants need to identify which of the three object classes was predictive of category membership for that rule (i.e., participants needed to solve for the rule dimension). Theories of learned selective attention posit that individuals will shift their attention away from feature dimensions which result in response errors, and through experience learn to focus on only information which is predictive of the outcome (e.g., Kruschke, 1992). Employing an independent localizer and task stimuli that are characterized by readily distinguishable distributed activation patterns in object-selective cortex (Haxby et al., 2001; O’Toole et al., 2005) enabled us to examine the operation of feature-based selective attention using representational similarity analysis (Kriegeskorte, Mur, & Bandettini, 2008). To conduct this analysis, the patterns participants activated during learning trials of the experimental task were compared to the patterns that were activated for face, object, and scene stimuli from the independent localizer task within a combined functional ROI that spanned face-, object-, and scene-selective cortical regions (Supplementary Figure A1). It was predicted that neural similarity to the subsequently chosen object class would increase over trials leading up to a decision, reflecting an attentional narrowing to dimensions that participants believed to be predictive (regardless of whether or not their belief was correct). Alternatively, we predicted that mean neural similarity to the non-rule/non-predictive dimensions would decrease as a function of time leading up to a decision.

The time course of neural similarities to both the object class participants subsequently chose as the rule dimension (similarity to chosen class) and the classes constituting the unchosen dimensions (similarity to unchosen class) is displayed in Figure 3. As hypothesized, there was a strong increasing relationship between learning duration over the five trials preceding a solve decision and neural similarity to the subsequently chosen item class (Figure 3, blue line), with mean similarity to chosen class increasing as subjects approached a decision (γ = .019, *t* = 3.34, *p* < .001). For similarity to the unchosen class (Figure 3, green line), the n-back to decision variable was also predictive of similarity output (γ = −.006, *t* = −2.00, *p* = .045), with similarity to unchosen items decreasing over the trials preceding decisions to stop learning. In both cases, the addition of a random slope parameter for n-back to decision did not improve model fit according to AIC comparison, suggesting no significant subject-level differences in the relationship between neural similarity to the object classes and trial number. The addition of random slopes did not affect the main statistical outcome of either model. Considering the two time-courses of neural similarity, it is important to note that the initial advantage of similarity output for the unchosen object classes (Figure 3) is attributable to the fact that early in learning, participants are more likely to be attending to a subsequently unchosen class due to their 2:1 frequency on each trial (see Figure 1).

**Figure 3.**
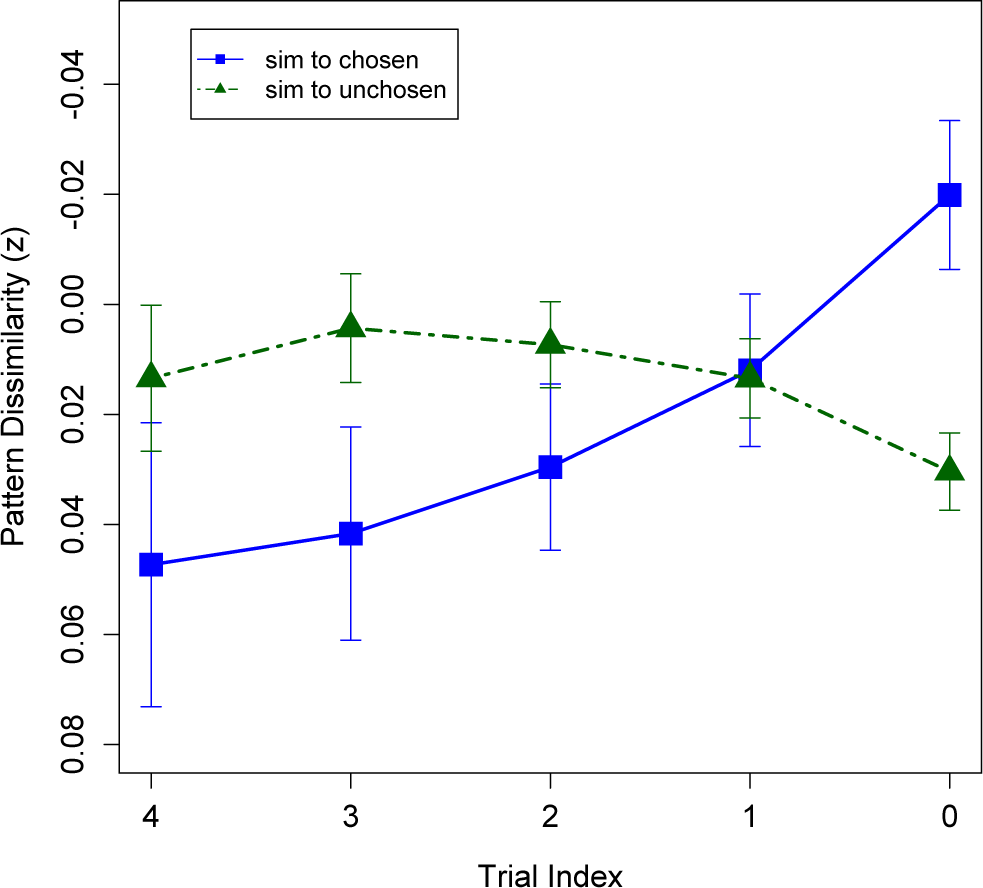
Mean time course of neural
similarities preceding solve decisions. On the x-axis, 4:0 represent category learning trials preceding a solve decision, with zero being the final learning trial before participants opted to stop and solve the rule. The y-axis displays z-scored mean multi-voxel pattern dissimilarity. Similarity estimates were computed in functionally-defined regions exhibiting selectivity for faces, objects, and scenes during the localizer phase (Supplementary Figure A1). The y-axis is flipped for ease of interpretation, as decreasing values signify increasing neural similarity. “Sim to chosen” (blue) reflects the neural similarity to the feature class that was eventually chosen as the category rule. “Sim to unchosen” (green) reflects the average neural similarity to the two feature classes that were not chosen as the category rule.

Due to the wide variability in rule solving accuracy observed in the present experiment, the overall patterns of neural similarity depicted in Figure 3 may mask differences in how the stimulus dimensions were attended across trials where learning was effective versus ineffective. Specifically, in light of the observed positive relationship between the average number of learning trials per rule and subject-level accuracy (Section 3.1), attention to chosen features may increase more gradually preceding correct, versus incorrect solves. Accordingly, we examined how neural similarity to chosen and unchosen rule dimensions varied during learning depending on whether the eventual rule selection following these trials was correct or incorrect. The time courses of neural similarity to chosen and unchosen object classes broken down by correct and incorrect solve responses are displayed in Figure 4 (correct solve; left panel, incorrect solve; right panel).

**Figure 4.**
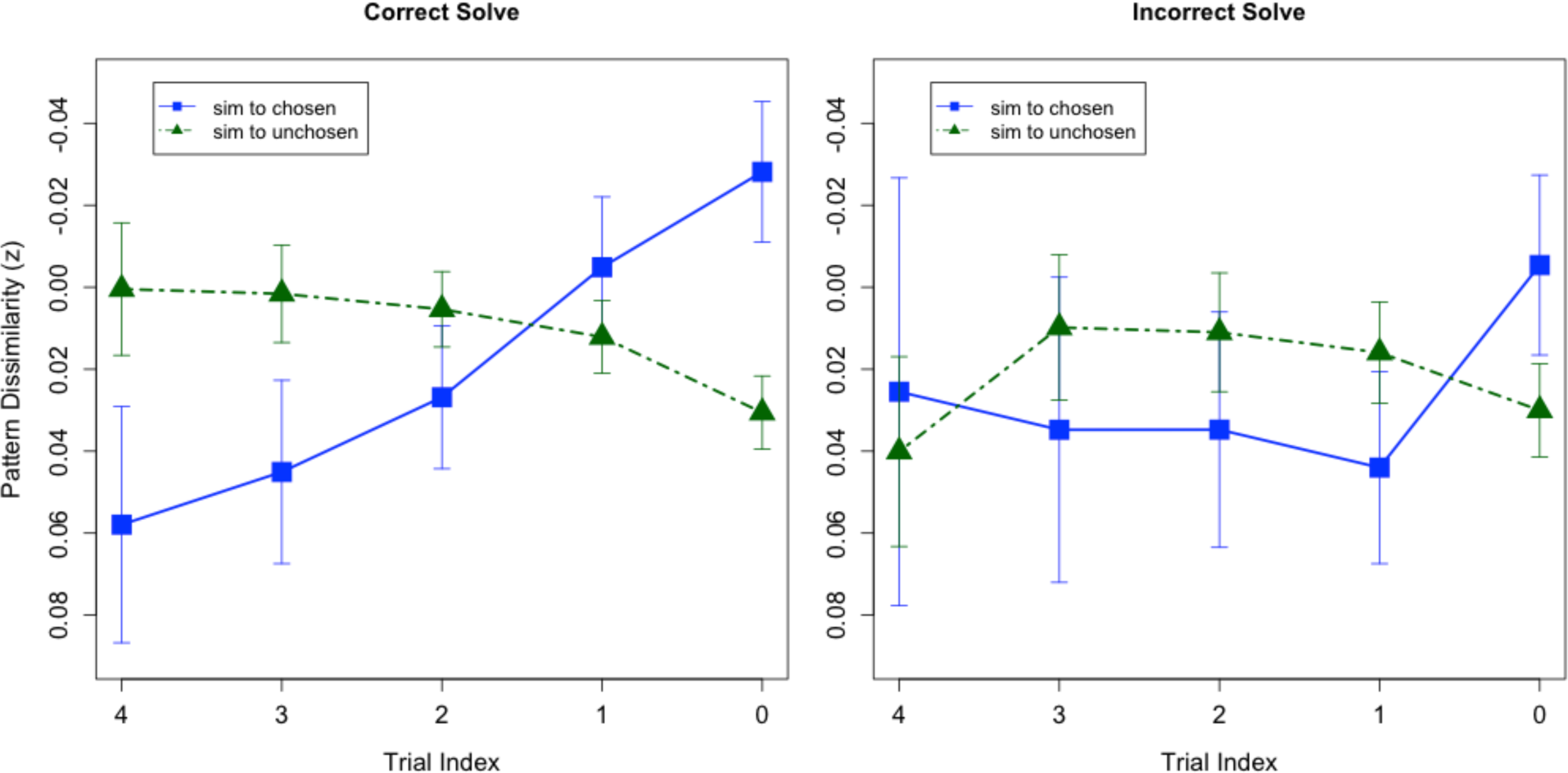
Mean time course of neural similarities to the chosen item class preceding correct versus incorrect solve decisions. On the x-axis, 4:0 represent category learning trials preceding a solve decision, with zero being the final learning trial before participants opted to stop and solve the rule. The y-axis displays *z*-scored mean multi-voxel pattern dissimilarity. Similarity estimates were computed in functionally-defined regions exhibiting selectivity for faces, objects, and scenes during the localizer phase (Supplementary Figure A1). The y-axis is flipped for ease of interpretation, as decreasing values signify increasing neural similarity. “Sim to chosen” (blue) reflects the neural similarity to the feature class that was eventually chosen as the category rule. “Sim to unchosen” (green) reflects the average neural similarity to the two feature classes that were not chosen as the category rule.

As with the combined analysis, we tested whether similarities to the chosen and unchosen dimensions exhibited significant linear effects over the five trials leading up to a solve decision. Prior to a correct response, we found a significant positive relationship between learning duration and neural similarity to the chosen object class (γ = .028, *t* = 4.18, *p* < .001), in addition to a significant negative relationship between learning duration and similarity to the unchosen object classes (γ = −.011, *t* = −3.15, *p* = .002). Incorrect responses were not preceded by linear changes in similarity over the most recent five learning trials for chosen (γ = .008, *t* = .814, *p* = .416) or unchosen (γ = −.001, *t* = −.239, *p* = .811) object classes. Consistent with the model combined across correct and incorrect trials, including random slope parameters for n-back was not merited based on AIC comparison. Taken together, these results suggest that when participants successfully solved the category rules, attention to the predictive object class increased more gradually leading up to a decision, accompanied by a similar decrement in attention to the two unpredictive object classes. Alternatively, no significant linear effects were observed for similarity to the chosen or unchosen object classes preceding incorrect solve decisions (Figure 4, right panel), suggesting that these choices may have been based on evidence accumulated over a shorter time course or possibly a single correct trial, consistent with the learning curve depicted in Figure 2.

We next sought to test whether trial-by-trial fluctuations in similarity to chosen class was predictive of stopping, using the binary decision to solve or continue immediately following each learning trial as the dependent variable. Indeed, a multi-level logistic regression revealed a significant positive relationship between similarity to chosen class on learning trials and the choice to solve the rule (γ = −.319, *z* = −2.99, *p* = .003), allowing each subject to have a random intercept. Including a random slope parameter for similarity to chosen class did not improve model fit based on AIC comparison, and the addition of the random parameter did not affect the statistical significance of this test. Additionally, we found a significant relationship between similarities to unchosen object classes and stopping decisions, with greater neural similarity to these stimulus dimensions predictive of decisions to continue learning (γ = .585, *z* = 2.84, *p* = .004). Again, adding a random slope parameter did not improve the model fit based on AIC, but its inclusion did not affect the statistical outcome of the test.

In sum, our RSA findings for the learning phase are consistent with the hypothesis that learned selective attention to a perceived rule dimension will be reflected by changes in the underlying neural similarity space, and that these attentionally-driven neural dynamics are predictive of stopping decisions in self-paced categorization. Moreover, differences in the time course of similarities to the chosen and unchosen object classes preceding correct, versus incorrect solves suggest that attentional changes over the course of learning may be strongly linked to participants’ learning strategies: correct solves were preceded by steady, linear increases in relative attention to the chosen feature, whereas incorrect solves were marked by greater relative attention to the chosen feature only on trials immediately preceding the solution.

### 3.3. Association Between Neural Similarity and Accuracy on Decision Trials

While the analysis of neural similarity to chosen and unchosen object classes during learning revealed trial-level differences in the attention to predictive cues for correct versus incorrect stopping decisions, another question of interest is whether overall task accuracy is related to the extent that participants activate the representations of predictive stimulus dimensions when considering whether to stop or continue learning. In contrast to alternative measures of visual attention such as eye tracking, a significant advantage of our MVPA measure is that the activation of each cue pattern may be compared to the relative activation of alternative cues on a single trial, regardless of whether the relevant stimuli are displayed on the screen at a given time (e.g., Zeithamova, Dominick, & Preston, 2012, Lewis-Peacock et al., 2012). We predicted that participants who were more accurate in their inferences would show a greater degree of neural similarity to the *correct* object class on decision trials, despite the fact that none of the stimuli were immediately available to the participants during these trials. To test this prediction, we computed a Pearson correlation between subject-level accuracy and mean similarity to the correct object class across decision trials in the task. As hypothesized, rule solving accuracy was significantly associated with greater neural similarity to the correct object class on decision trials (*r* = −.510, *t* (23) = −2.84, *p* = .009), suggesting that individuals who performed more accurately on the task activated patterns associated with the predictive visual stimulus dimension to a greater extent than those who performed less accurately. The correlation between subject-level accuracy and similarity to the correct object class is depicted in Figure 5.

**Figure 5.**
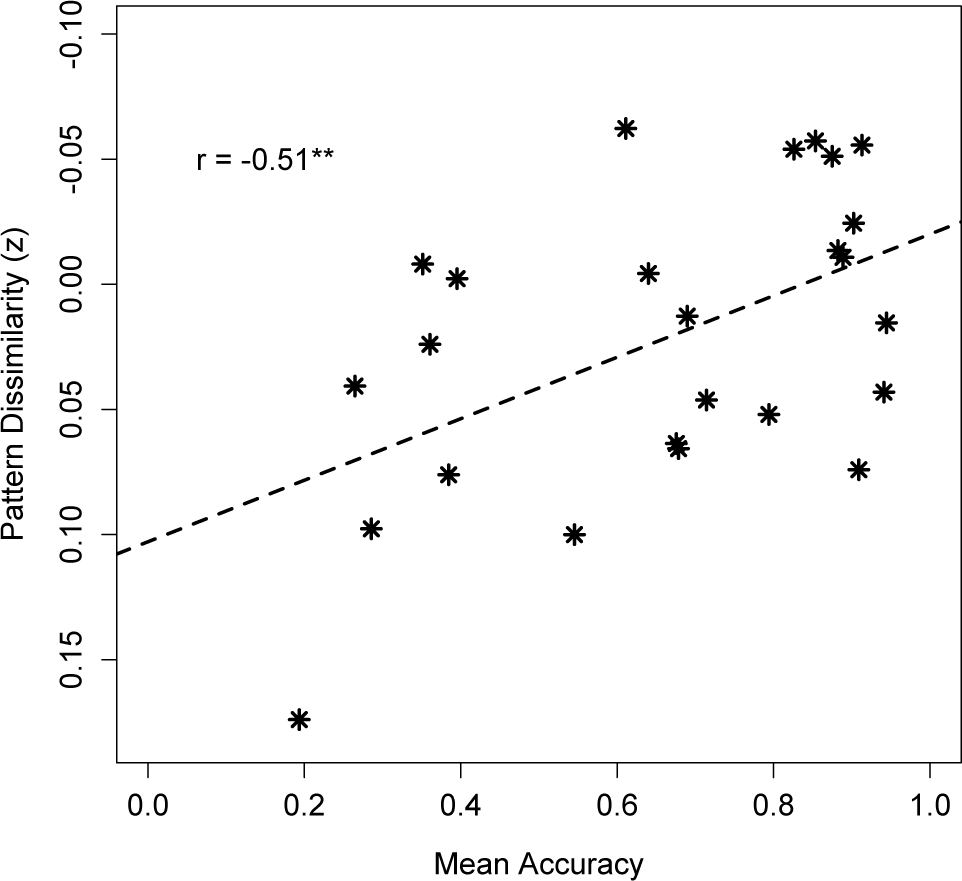
Association between subject-level rule solving accuracy and similarity to the correct object class on decision trials. The y-axis displays zscored mean multi-voxel pattern dissimilarity in ventral occipitotemporal cortex (Supplementary Figure A1). The y-axis is flipped for ease of interpretation, as decreasing values signify increasing neural similarity.

### 3.4. Time-course of vmPFC Signal

Based on recent findings suggesting that the vmPFC tracks relative decision evidence in categorization (Davis, Goldwater, & Giron, 2017) and subjective confidence associated value-based decisions (Di Martino et al., 2013; Barron, Garvert, & Behrens, 2015; Lebreton et al., 2015), we hypothesized that activation in this region would be associated with stopping decisions in self-paced learning. A series of task-relevant questions pertaining to this prediction are how signal changes in vmPFC might covary with attention to relevant feature dimensions, the temporal distance from a solve decision, and individual differences in rule solving accuracy. In light of the strong relationship between neural similarity to chosen/unchosen object classes during learning and decisions to solve the rule, we first aimed to test whether similarity patterns during learning were predictive of vmPFC signal on the immediately following decision trial, and whether this relationship may be moderated by the temporal distance from a solve decision and subject-level accuracy. We thus computed a multi-level model over decision trials with trial-level vmPFC signal as the outcome variable, and learning similarities to the chosen/unchosen object classes, 5-trial n-back, and subject-level accuracy as predictor variables. The model tested three-way interactions between similarity to the chosen/unchosen object classes, n-back, and accuracy. Interaction effects related to vmPFC signal at the time of decision were not statistically significant, while main effects for similarity to the chosen (γ = −.981, *t* = −1.92, *p* = .056) and unchosen (γ = −1.92, *t* = −1.92, *p* = .055) object classes during learning were found to be marginally significant. Given the direction of the relationships between similarity to chosen/unchosen object classes during learning and vmPFC signal on decision trials, our results suggest that vmPFC may be responsive to how strongly individual cues are attended during learning, irrespective of whether the cue is eventually chosen as the relevant rule dimension.

In Section 3.3, we show that participants’ overall accuracy is strongly associated with reactivation of correct cue patterns on decision trials. A related question of interest is how reactivation of chosen/unchosen object classes during the decision phase may relate to trial-level activation of vmPFC. Accordingly, we conducted a multi-level regression with vmPFC signal as the outcome variable, using similarities to the chosen/unchosen object classes on decision trials, 5-trial n-back, and subject-level accuracy as predictor variables. Three-way interactions among similarity to the chosen/unchosen object classes, n-back, and accuracy were tested. Results of the multi-level model revealed a significant two-way interaction between n-back and accuracy in predicting vmPFC signal (γ = −.154, *t* = −2.45, *p* = .014). Additionally, the main effect of similarity to the unchosen object class on decision trials was marginally significant (γ = − 1.79, *t* = −1.96, *p* = .05). The interpretation of the interaction effect between n-back and accuracy is that a stronger relationship exists between temporal distance from a solve decision and vmPFC signal among higher, versus lower accuracy participants: more accurate participants exhibited a steeper slope of vmPFC activation over time relative to less accurate participants. For illustrative purposes, this interaction effect is depicted in Figure 6 for the top and bottom 1/3 of task performers (all subjects were included in the statistical test).

**Figure 6.**
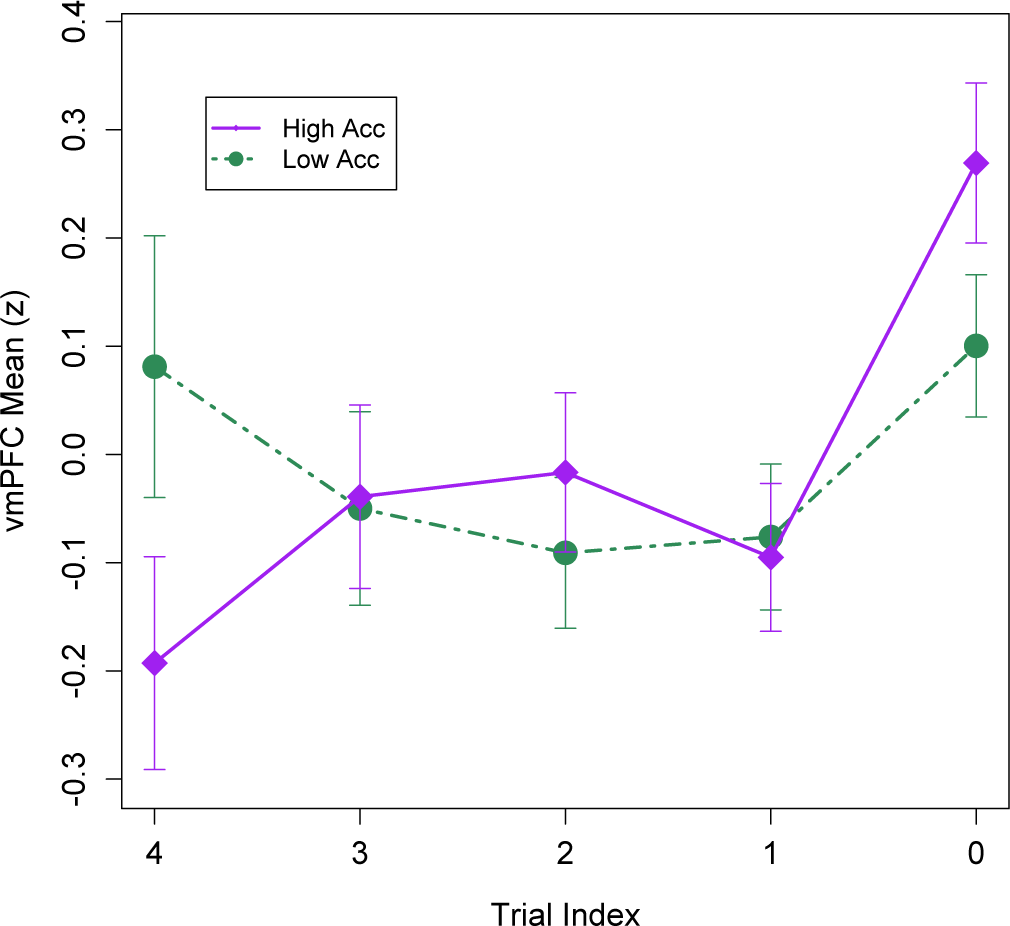
Mean trial-level BOLD
response in vmPFC for accurate and inaccurate participants. On the x-axis, 4:0 represent the decision trials preceding a solve decision, with zero being the decision trial where participants opted to stop and solve the rule. The y-axis displays z-scored mean BOLD activation in anatomically-defined vmPFC as measured by the betaseries regression. “High Acc” (purple) corresponds to the top 1/3 of task performers (n = 8, *M* = 90.7% correct), while “Low Acc” (green) corresponds to the bottom 1/3 of task performers (n = 8, *M* = 34.8% correct).

Similar to the binary decision model based on learning similarities (Section 3.2), we tested whether trial-by-trial fluctuations in vmPFC signal during decision trials was predictive of stopping, where the dependent variable was whether the participant chose to solve the rule or continue learning on a given trial. Consistent with our findings for the RSA attention measure, a multi-level logistic regression revealed a significant positive relationship between vmPFC signal on decision trials and whether or not subjects chose to stop (γ = 0.274, *z* = 2.76, *p* = .006), allowing each subject to have a random intercept. The inclusion of a random slope parameter for the relationship between vmPFC signal and stopping was found to be merited based on AIC comparison between model fits, suggesting significant subject-level variability in the relationship between vmPFC activation and stopping at the trial level.

### 3.5. Neural Correlates of Stopping vs. Continuing on Decision Trials

It was hypothesized that decisions to stop learning and solve the rules would be more cognitively demanding than choices to continue learning, and would engage a network including prefrontal cortex, basal ganglia, and visual association cortices. Specifically, we expected that ventral occipitotemporal cortex would be activated because this region is associated with direct sensory or memory-based evidence for an object class (O’Craven & Kanwisher, 2000; Haxby et al., 2001), and that vmPFC would be engaged because of its association with high relative confidence and overall evidence for a decision (De Martino et al., 2013; Barron, Garvert, & Behrens, 2015; Lebreton et al., 2015; Davis, Goldwater, & Giron, 2017). We contrasted Solve > Continue on decision trials, finding a single, widespread cluster of activation (Figure 7A; Table 1). Within the cluster, the peak BOLD activity was located in medial PFC, spanning bilateral paracingulate gyrus, vmPFC, anterior cingulate cortex (ACC), and frontal pole. Additional regions exhibiting significant activation for Solve > Continue included (but were not limited to) bilateral ventral occipitotemporal cortex, anterior insula, and ventral striatum. No regions were significantly activated for the contrast Continue > Solve. Our findings support the prediction that stopping an information search, compared with decisions to continue learning, engages a broad network including regions that represent visual category information and accumulated decision evidence. Moreover, the co-activation of dorsal ACC, ventral striatum, and insula within this contrast suggests that self-initiated stopping decisions involve a similar network of brain regions to those engaged for rule switching in matching and classification tasks (Simard et al., 2011; Liu et al., 2015). All final statistical maps are available at <u>https://osf.io/ju9mz/</u> for further exploration.

**Figure 7.**
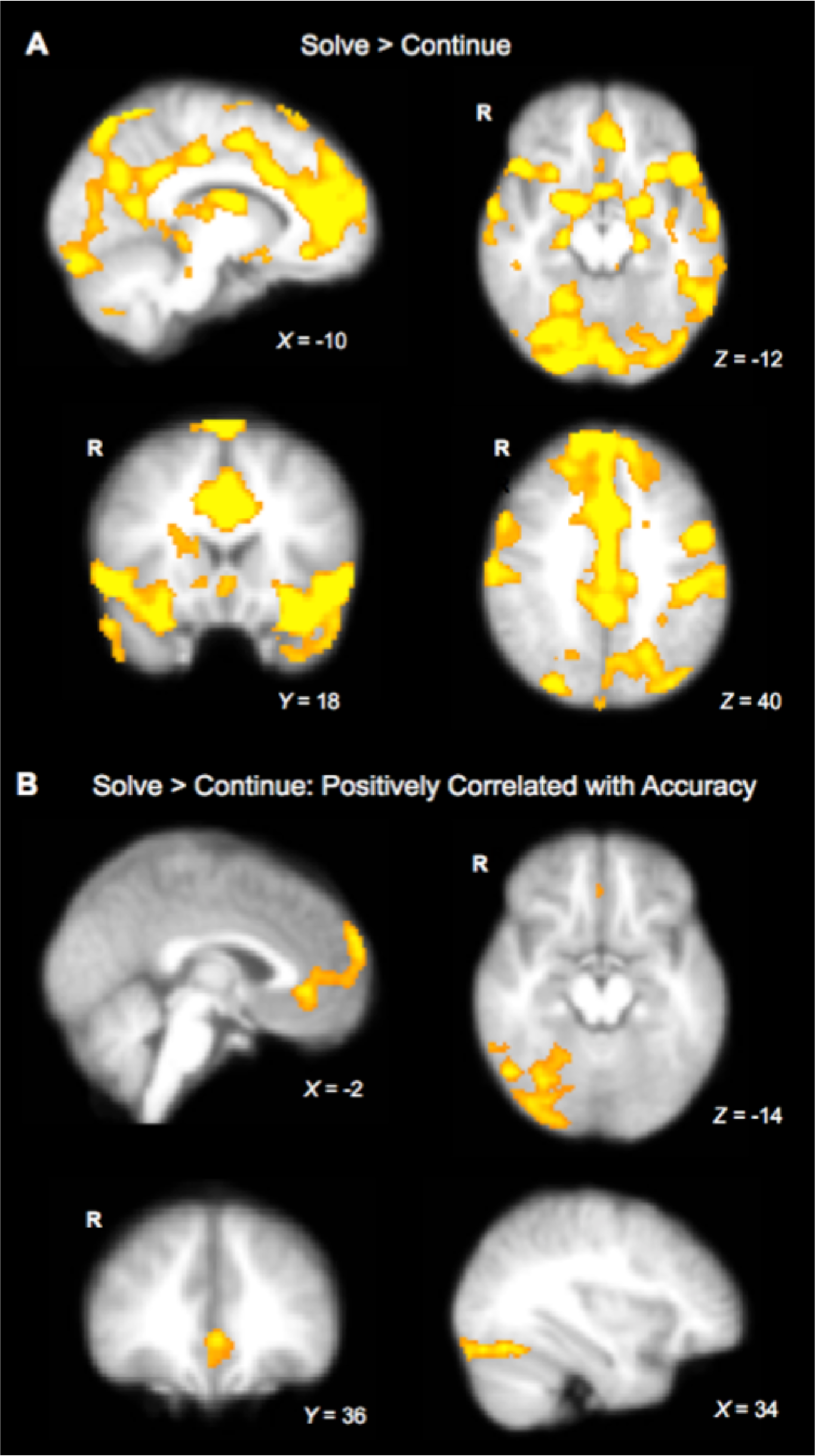
Univariate fMRI results. A) Brain regions activated for Solve > Continue on decision trials. B) Regions where activation was positively correlated with mean solve accuracy within the Solve > Continue contrast.

### 3.6. Moderation of Stopping vs. Continuing Activation by Solving Accuracy

As stopping, in the current task, is an inherently self-governed decision, it was predicted that regional engagement on decision trials would differ between subjects as a function of performance. Specifically, we examined mean rule solving accuracy as a subject-level moderator for the contrast Solve > Continue. It was predicted that individuals who were more accurate in rule solving would exhibit more sensitivity in the vmPFC and regions that discriminate between faces, objects, and scenes. Two clusters were positively correlated with accuracy on stopping trials (Figure 7B; Table 1). The first cluster was located in right ventral visual cortex, with maxima in lateral occipital complex and occipital fusiform gyrus. The second included medial frontal pole, rostral anterior cingulate, and vmPFC. The engagement of our *a priori* ROIs in the moderation suggests that they are involved in *successful* stopping, as their activity during stop decisions is correlated with the likelihood of subjects knowing the correct rule. Consistent with the prediction that successful stopping decisions would require learned selective attention to predictive object classes, the activation of category selective regions suggests that rule solving accuracy is supported by greater relative consideration of the features associated with a rule when deciding whether to stop. In addition to our attentional predictions, we hypothesized that vmPFC would be associated with task accuracy, as high-performing participants are likely to accumulate more evidence for their decisions and, consequently, be more confident at the time of inference. The observed relationship between vmPFC engagement and task performance confirmed this prediction, and is consistent with recent findings implicating vmPFC in representing high relative decision evidence (Davis, Goldwater, & Giron, 2017) and the confidence associated with value judgments (De Martino et al., 2013; Barron, Garvert, & Behrens, 2015; Lebreton et al., 2015) in decision making.

### 3.7. PPI Results

A crucial question concerning the neurobiological basis of stopping an information search is whether there is a specific region responsible for integrating evidence from accumulator regions to precipitate a decision. To address this question, we conducted a PPI analysis to determine where the BOLD signal covaried with activation in vmPFC (Figure 8A) on stopping trials. Based on previous research illustrating its role as an integrator region that responds to inputs from vmPFC (Baumgartner et al., 2011; Rudorf & Hare, 2014), we predicted that dlPFC would play a critical role in the execution of stopping once a threshold is reached. Consistent with this hypothesis, the PPI analysis revealed activation in bilateral dlPFC that was functionally coupled with the vmPFC seed for Solve > Continue on decision trials (Figure 8B). In addition to dlPFC, signal in dorsomedial PFC, pre-SMA, and frontal pole displayed enhanced functional connectivity with vmPFC when participants chose to solve. With respect to the frontal pole, we specifically found that the primarily dorsal/lateral PFC cluster extended rostrally to a portion of right rostrolateral PFC (rlPFC) previously associated with metacognitive accuracy in value-based choice (Di Martino et al., 2013). This region often tracks rule-evaluation processes when tasks encourage use of higher-level symbolic representations (Badre, Kayser, & D’Esposito, 2010; Davis, Goldwater, & Giron, 2017; Paniukov & Davis, 2017).

**Figure 8.**
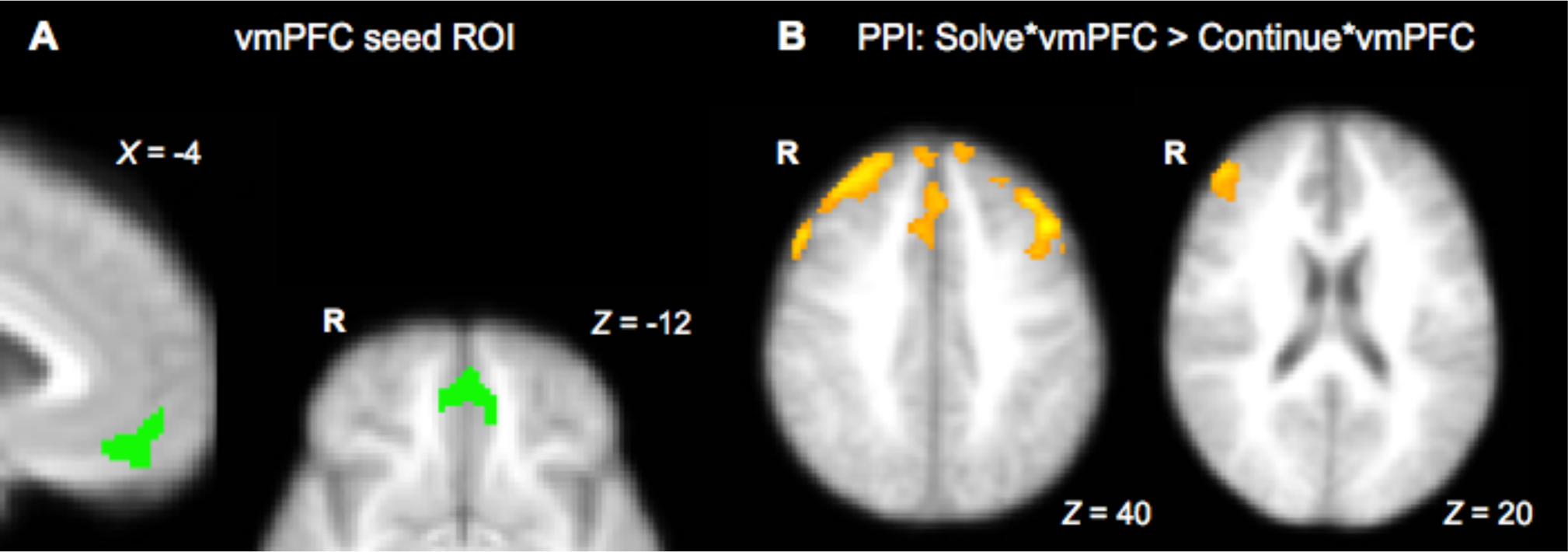
Results of the PPI analysis. A) The vmPFC seed region used in the PPI. B) Regions where neural activation covaried with the BOLD time series in vmPFC on decision trials receiving a solve response versus a continue response.

## 4. Discussion

In real-world decision environments, learners are responsible for regulating the amount of time they spend gathering information about a concept before making a choice. This regulation process is instantiated by people’s decisions to stop learning. To understand the neural mechanisms responsible for stopping of information search in humans, we used fMRI and a self-paced category learning task where participants learned unidimensional rules where a single visual feature was predictive of category membership. Our fMRI results show that compared with continuing to learn, decisions to stop engage a broad network of brain regions including PFC, basal ganglia, and visual association cortices. Behaviorally, the sample varied in desired learning length (time-to-stop) and rule-solving accuracy. Higher task accuracy was predictive of greater activation in vmPFC, frontal pole, and right occipitotemporal cortex on stopping trials. Using MVPA, we tracked neural similarity to the features (face, object, or scene) during learning that were eventually chosen to be the predictive class. This analysis revealed a significant linear increase of similarity to chosen classes over the five learning trials prior to a solve response, and further, that pattern similarities were predictive of choice on a trial-by-trial basis. Finally, a psychophysiological interaction analysis revealed that the dlPFC was more functionally coupled with the vmPFC when subjects chose to stop versus continue, consistent with its hypothesized role in monitoring accumulated evidence to determine whether a decision threshold has been reached.

In the context of our rule-based, multi-feature category learning task, individuals must learn to distinguish relevant from irrelevant information to accurately classify exemplars. Rather than measuring fluctuations in mean activation within our ventral visual ROI in response to varying task conditions, the use of MVPA allowed us to estimate the extent to which participants were focusing on each object class over the course of the task. Our analysis revealed that selective attention to object classes that subjects eventually chose as the rule dimension when solving a rule displayed an approximately linear ramping effect over the trials preceding correct solve decisions. Further, trial-by-trial estimates of representational similarity to chosen classes during learning were predictive of decisions to stop learning and solve the rule immediately following the trial. These findings extend previous research suggesting that neural representations of stimuli are modified over the course of category learning to facilitate optimal performance within a variety of task structures (Mack, Love, & Preston, 2013; Folstein, Palmeri, & Gauthier, 2013; Davis & Poldrack, 2014) by showing how learning-dependent changes in the neural similarity space relate to subjective decision criteria. To our knowledge, this study is the first to examine the time course of dimensional selective attention in self-paced learning.

The vmPFC is a region that we hypothesized would be more activated for stopping versus the choice to continue learning. Past research has thoroughly established a role of the vmPFC in reward prediction and subjective value assessment (Tom et al., 2007; Bartra et al., 2013). Due to the absence of extrinsic rewards for correct responses or differing outcomes that were mapped on to each feature, in this study it was expected that the vmPFC would serve a more general function of tracking evidence for category rules (Davis, Goldwater, & Giron, 2017). Indeed, we found vmPFC to be one of the regions involved in stopping, and that activation in vmPFC covaried with task accuracy at the time subjects chose to stop. These results lend support to an emerging area of research that has implicated vmPFC in representing subjective confidence in parallel with value (De Martino et al., 2013; Barron, Garvert, & Behrens, 2015; Lebreton et al., 2015). It is important to note that the present study is unable to dissociate the neural signals associated with confidence from those associated with implicit value, given that the goal of correctly solving as many rules as possible necessarily correlates participants’ confidence with reward maximization. Accordingly, future studies may seek to address whether the vmPFC is particularly sensitive to confidence, value, or a combination of these factors in self-paced learning, perhaps by manipulating the reward associated with different rule dimensions. However, the present results suggest that in learning paradigms that demand varying attention to different visual features, accumulation processes in vmPFC are closely linked to absolute differences in the strength of competing feature representations. The possibility that vmPFC tracks the strength of signal in relevant perceptual regions when weighing a decision accords with our finding that greater task accuracy was associated with higher relative activation in both vmPFC and ventral occipitotemporal cortex. From a learned selective attention standpoint, individuals who represent predictive features the most strongly should both be more confident in their decisions and more accurate in their inferences.

Our results extend recent research examining the neural correlates of decision evidence in categorization. Using a temporally extended classification task wherein probabilistic cues were presented to participants incrementally within a trial, Braunlich and Seger (2016) employed a model-based fMRI approach to isolate brain regions associated with parametric modulations in decision evidence that were independent of response‐ and urgency-related signals over time. Their results revealed that the absolute value of decision evidence accumulated over a trial correlated with activation in rostral and dorsolateral PFC, but surprisingly, was not associated with increased signal in vmPFC. One possibility for the divergence in results between this study and the current experiment regarding vmPFC is that this region is specifically engaged in relation to subjective appraisals of decision evidence, as opposed to reflecting the objective amount of evidence a person is exposed to. The regions engaged for stopping decisions in our study are not inherently tied to the objective evidence for a rule, as reflected in the high degree of variability in the average number of trials participants chose to view before opting to solve the rules. Rather, by their very nature, these stopping decisions are a reflection of participants’ subjective estimates of making a correct response. This idea accords with findings in the memory literature that implicate vmPFC in tracking metacognitive “feeling-of-knowing” judgments associated with retrieval attempts, but not retrieval accuracy itself (Schnyer, Nichols, & Verfaellie, 2005). Taken together, we suggest that while rostral and dorsal subregions of the PFC appear to be critical for performing computations related to finite differences in decision evidence (e.g., Heekeren et al., 2004; Braunlich & Seger, 2016; Davis, Goldwater & Giron, 2017), vmPFC may uniquely represent more subjective decision variables such as confidence and the perceived costs and benefits associated with a decision (e.g., Basten et al., 2010).

Through a psychophysiological interaction analysis, we found that activation in vmPFC during stopping decisions is coupled with modulation of dlPFC. These findings are generally consistent with previous research showing that dlPFC interacts dynamically with vmPFC by comparing value signals represented in the latter region, and that dlPFC may precipitate responses based on this comparison process (Pochon et al., 2002; Baumgartner et al., 2011; Kahnt et al., 2011; Rudorf & Hare, 2014). When subjects chose to stop learning and respond to the rule, activity in vmPFC covaried with dlPFC to a greater extent than when they were not yet ready to solve. Accordingly, dlPFC may compute whether a decision threshold has been reached based on the relative evidence that has accumulated for a given category rule and participants’ meta-knowledge of the task structure. Our results regarding vmPFC-dlPFC interactions in self-paced category learning serve as a bridge between research focused on simple perceptual decisions and studies that primarily focus on the expected reward or monetary gains/losses incurred from a decision, and suggest that a common neural mechanism may support stopping in both category learning and value-based decision making. Because of the relatively indirect connection between dlPFC activity and stopping behavior in the present study, future research employing model-based fMRI may be useful for testing explicit predictions about the putative executive role of dlPFC in self-paced learning.

As an alternative to stopping per se, decisions to solve the category rule are involve a switching process, whereby participants explicitly transition from hypothesis testing to solving the perceived category rule. Accordingly, contextualizing our findings in light of the extensive hypothesis testing/rule switching literature may be useful for understanding the various strategies participants may have employed to navigate our stopping task. Neurally, a subset of the regions engaged for Solve > Continue are consistent with those often implicated in rule switching, including the ventral striatum (Seger & Cincotta, 2006; Simard et al., 2011; Liu et al., 2015) and regions that comprise the salience network (dorsal anterior cingulate, anterior insula, and inferior frontal cortex; Dosenbach et al., 2006; Sridharan, Leviton, & Menon, 2008; Liu et al., 2015). Behaviorally, while an optimal observer may adopt a win-stay lose-shift strategy in matching tasks where correct feedback is fully correlated with knowledge of the correct rule, learners often rely on this strategy in multidimensional classification tasks despite the fact that a string of correct responses may fail to eliminate viable alternative rules (Shepard, Hovland, & Jenkins, 1961, Nosofsky, Palmeri, & McKinley, 1994, Niv et al., 2015; Paniukov & Davis, 2017). We found subject-level rule solving accuracy to be strongly associated with the number of learning trials participants encountered before choosing to solve the rule, with less accurate participants choosing to solve after fewer learning trials and fewer instances of correct feedback than those with high accuracy. As such, it is possible that poorer task performers in the current study frequently adopted a suboptimal win-stay lose-shift strategy when deciding whether they were prepared to solve the rule, in some cases using a single correct trial as justification to move away from hypothesis testing and respond with the rule. Our attentional tuning results mirror this possibility, as incorrect solves were preceded by a marked increase in neural similarity to the chosen object class on the final learning trial, as opposed to correct solves where similarity to the chosen class increased in a more gradual manner (Figure 4).

### 4.1. Conclusions

To balance accuracy and efficiency in decision making, learners must determine when their current knowledge state matches the knowledge that is required to achieve a particular goal. Using representational similarity analysis, we illustrate that such stopping decisions are preceded by greater activation of patterns consistent with perceived rule dimensions, consistent with theories of learned selective attention. Although stopping engaged a widely distributed network in the brain, our findings suggest that vmPFC and object-selective cortical regions are differentially engaged according to participants’ tendencies to accurately solve rules, and thus these regions may play unique roles in supporting accurate and timely stopping decisions in self-paced category learning. Finally, our results demonstrate that dlPFC is functionally coupled with vmPFC during stopping decisions, suggesting that this region may serve a similar criterion-setting function in self-paced learning to that observed across studies of perceptual and value-based decision making.

## AUTHOR CONTRIBUTIONS

SO, EW, MS, and TD designed the study and wrote the paper. SO and TD analyzed the fMRI data. SO collected the fMRI data for the experiment.

## ACKNOWLEDGEMENTS

Funding: This work was supported by an internal grant awarded to TD and EW from Texas Tech University.

